# Cure of congenital purpura fulminans via expression of engineered protein C through neonatal genome editing in mice

**DOI:** 10.1101/2023.02.26.530058

**Authors:** Tomoki Togashi, Nemekhbayar Baatartsogt, Yasumitsu Nagao, Yuji Kashiwakura, Morisada Hayakawa, Nobuhiko Kamoshita, Takafumi Hiramoto, Takayuki Fujiwara, Eriko Morishita, Osamu Nureki, Tsukasa Ohmori

## Abstract

Protein C (PC) is a plasma anticoagulant encoded by *PROC*; mutation in both *PROC* alleles results in neonatal purpura fulminans—a fatal systemic thrombotic disorder. In the present study, we aimed to develop a genome editing treatment to cure congenital PC deficiency. First, we generated an engineered activated PC to insert a self-cleaving peptide sequence between light and heavy chains. The engineered PC could be released in its activated form and significantly prolonged the plasma coagulation time independent of the cofactor activity of protein S *in vitro*. The adeno-associated virus (AAV) vector-mediated expression of the engineered PC, but not wild-type PC, prolonged coagulation time owing to the inhibition of activated coagulation factor V in a dose-dependent manner and abolished pathological thrombus formation *in vivo* in C57BL/6 mice. The insertion of *EGFP* sequence conjugated with self-cleaving peptide sequence at *Alb* locus via neonatal *in vivo* genome editing using AAV vector resulted in the expression of EGFP in 7% of liver cells, mainly via homology-directed repair, in mice. Finally, we succeeded in improving the survival of PC-deficient mice by expressing the engineered PC via neonatal genome editing *in vivo*. These results suggest that the expression of the engineered PC via neonatal genome editing is a potential cure for severe congenital PC deficiency.

**One Sentence Summary:** Ectopic expression of an engineered protein C via genome editing cures protein C deficiency in mice.

## INTRODUCTION

Thromboembolic disorders account for one in four deaths and are the leading cause of mortality worldwide *(1)*. Thrombogenesis or thrombus formation in the blood is determined by the balance between the activities of coagulation and anticoagulation factors *(2)*. The decreased activity of anticoagulation factors, including antithrombin (AT), protein C (PC), and protein S (PS), in the blood is an important predisposing factor for thrombogenicity *(3)*. AT directly inhibits activated coagulation factors including thrombin and coagulation factor X *(4)*, whereas PC circulates as a zymogen in the blood and is activated at the site of thrombus formation *(5, 6)*. Thrombin generated via the coagulation factor cascade binds to thrombomodulin on the vascular endothelial cells, thereby leading to the activation of PC via proteolytic cleavage *(5, 6)*. The activated protein C (APC) along with the cofactor PS degrades activated coagulation factor V (FV) and factor VIII (FVIII), thereby inhibiting thrombus propagation *(5, 6)*.

Genetic abnormality is a major cause of decreased levels of anticoagulation factors in the blood. Heterozygous abnormality of PC (*PROC*), PS (*PROS1*), and antithrombin (*SERPINC1*) genes is a well-known risk factor for venous thromboembolism in adulthood *(7)*. Abnormalities in both alleles of *PROC* or *PROS1* cause neonatal purpura fulminans, which is characterized by life-threatening systemic thrombosis and hemorrhagic skin necrosis shortly after birth *(8, 9)*. Although a replacement therapy using PC preparations is applied in the acute phase, it is not a realistic treatment approach that can be used throughout life because of the extremely short half-life of the protein *(10)*. Anticoagulation therapy with warfarin or heparin can be an alternative to inhibit thrombus formation for life *(11, 12)*. However, the patients remain at a risk of bleeding caused by extensive anticoagulation and thrombosis owing to the underlying disease throughout their lives *(13)*. Developing innovative therapies for PC or PS deficiency is required, particularly for patients who are homozygotes and combined heterozygotes with severe disease.

Recently, gene therapy and genome editing therapies are attracting attention as novel modalities for curing intractable diseases. Because PC is a protein produced in the liver hepatocytes in a manner similar to the production of coagulation factors, gene therapy and genome editing that are currently being developed for hemophilia B (coagulation factor IX [FIX] deficiency) may apply to PC deficiency. The plasma level of PC in human was reportedly 63 nmol/L *(10)*, which is comparable to that of FIX (87 nmol/L) *(14)*. Gene therapy for hemophilia B using a hyperactive mutant called the FIX Padua (R338L mutation) has shown effectiveness *(15, 16)*. PC is a serine protease that is primarily present as a zymogen in the blood. The concentration of APC in the blood is approximately 40 pmol/L, representing only 1/1,700 of the total PC *(17)*. We hypothesized that gene therapy or genome editing therapy can become a realistic alternative treatment approach if the active form of PC is secreted. We aimed to develop an engineered activated PC and assessed its application as gene therapy and genome editing therapy targeting PC deficiency in mice.

## RESULTS

### Prolongation of coagulation time by the engineered PC

We attempted to design an engineered human PC (hPC) to release the activated form. We inserted the furin cleavage sequences (KR, RKR, KRRKR, 2RKR, 3RKR, 4RKR, RHQR, or RSKR) or porcine teschorivus-1–derived sequence (P2A) into the thrombin recognition site of the *PROC* cDNAs (Fig. 1A). Following this, we generated HEK293 cells stably expressing hPC and measured hPC activity (hPC:C) in the supernatants. To compare the anticoagulant ability of engineered hPCs, we used the previously reported hPC with enhanced activity (PC-QGNSEDY: H10Q, S11G, S12N, E23S, N32E, N33D, and H44Y in light chain) *(18)* as a control. hPC:C was detected in the supernatants of cells expressing wild-type (WT) hPC (hPC-WT), hPC-QGNSEDY, and several engineered PCs (Fig. 1B). However, hPC:C was not detected in the supernatant of HEK293 cells expressing hPC-RKR, hPC-RHQR, and hPC-P2A (Fig. 1B). hPC:C is determined via the cleavage of a synthetic substrate by hPC, which is activated by snake venom in the reagent *(19)*. To verify whether our engineered hPCs were secreted in their active form, we measured hPC:C production in the absence of snake venom. Although hPC:C in the supernatants of cells expressing hPC-WT or hPC-QGNSEDY was not measured without snake venom, we detected a significant increase in hPC:C in the supernatants of cells expressing hPC-KRRKR, hPC-2RKR, hPC-3RKR, and hPC-4RKR (Fig. 1C). We selected hPC-2RKR for the subsequent experiments because of the highest secretion efficiency (Fig. 1B, C).

**Fig. 1.**
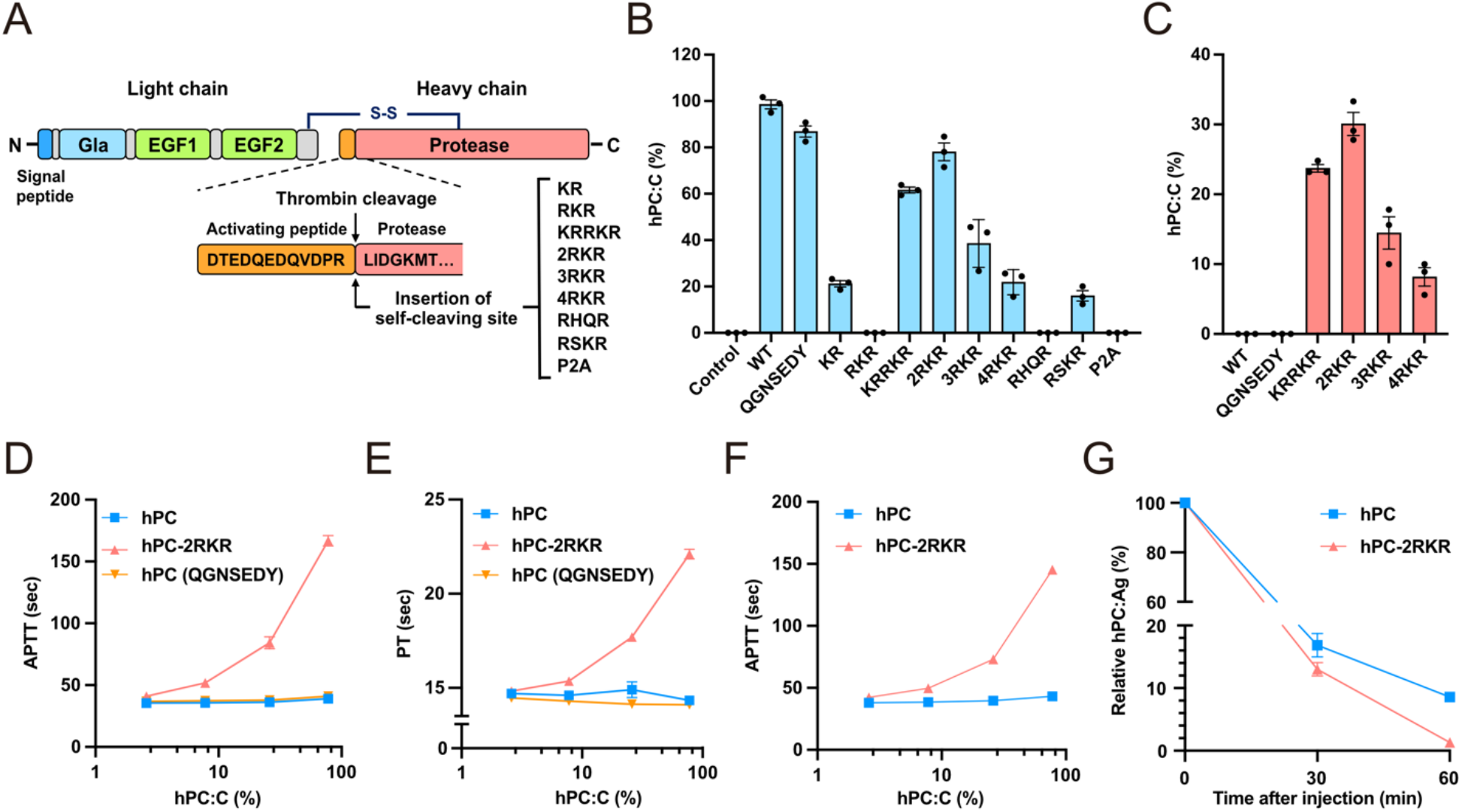
Generation of engineered activated hPC. (**A**) Schematic presentation of the engineered hPC. An indicated self-cleaving peptide sequence was inserted into the thrombin cleavage site of hPC. (**B**) hPC:C levels (mean ± SEM [n = 3–4]) in the supernatant obtained from the HEK293 cells stably expressing an indicated engineered hPC. (**C**) hPC:C levels (mean ± SEM [n = 3–4]) measured without snake venom. (**D, E**) Normal human plasma was incubated with an indicated concentration of wild-type hPC, the previously reported engineered hPC (QGNSEDY), or the engineered activated hPC (hPC-2RKR) for 15 min. APTT (**D**) and PT (**E**) (mean ± SEM [n = 3]) were measured using an automated coagulation analyzer. (**F**) Protein S-deficient plasma was incubated with an indicated concentration of hPC or hPC-2RKR for 15 min. APTT (mean ± SEM [n = 3]) was measured using an automated coagulation analyzer. (**G**) hPC or hPC-2RKR proteins obtained from the supernatant were intravenously injected into C57BL/6 mice. The blood was drawn at 30 and 60 min post-injection. Plasma levels of hPC antigen (hPC:Ag) (mean ± SEM [n = 3]) after the administration were measured using ELISA and expressed as a percentage immediately after administration. hPC, human protein C; hPC:C, human protein C activity; QGNSEDY, mutations at H10Q, S11G, S12N, E23S, N32E, N33D, and H44Y in hPC; APTT, activated partial thromboplastin time; PT, prothrombin time; hPC:Ag, protein C antigen.

We examined the anticoagulant potential of hPC-2RKR to measure the inhibition of human plasma coagulation. Coagulation time (activated partial thromboplastin time [APTT] and prothrombin time [PT]) of normal human plasma was assessed by mixing with an indicated concentration of an hPC or hPC-2RKR from the supernatant. hPC-WT and hPC-QGNSEDY did not affect APTT and PT, whereas hPC-2RKR significantly prolonged PT and APTT in a dose-dependent manner (Fig. 1D, E). Moreover, the prolongation of APTT and PT by hPC-2RKR was observed in PS-deficient plasma, suggesting that the anticoagulant activity of hPC-2RKR did not require the cofactor activity of PS (Fig. 1F). Because activated hPC has an extremely short half-life in plasma *(10)*, we compared the clearance of recombinant hPC-WT and hPC-2RKR in mice following an intravenous injection. As expected, compared with hPC-WT, hPC-2RKR exhibited more rapid clearance (Fig. 1G).

### Inhibition of pathogenic thrombus by the engineered PC *in vivo*

To assess the anticoagulant activity of PC-2RKR *in vivo*, we employed a mouse model of thrombus formation induced by reactive oxygen species (ROS) production; thrombus formation in mice was observed using intravital microscopy. We created an adeno-associated virus vector 8 (AAV8) harboring WT mouse PC (mPC) or the engineered mPC corresponding to hPC-2RKR (mPC-2RKR) and injected the vector in adult C57BL/6 mice at different doses (low dose, 4.0 × 10^10^ vg; medium dose, 1.2 × 10^11^ vg; and high dose, 4.0 × 10^11^ vg). In mice treated with the AAV8 vector harboring mPC (WT), although the mPC antigen (mPC:Ag) levels significantly increased in a dose-dependent manner, plasma coagulation time assessed by APTT remained unaffected even at a high dose (Fig. 2A, B). By contrast, in mice treated with the AAV8 vector harboring mPC-2RKR, coagulation time was prolonged in a dose-dependent manner; however, compared with mice expressing mPC (WT), those expressing mPC-2RKR showed lower increases in mPC:Ag levels (Fig. 2A, B). Furthermore, in mice expressing mPC-2RKR, a significant decrease in FV activity (FV:C) in plasma was observed (Fig. 2C). FVIII activity (FVIII:C) tended to decrease in a dose-dependent manner in mice treated with mPC-2RKR; however, compared with FV activity, mPC-2RKR expression exerted a lower effect on FVIII activity (Fig. 2D). Furthermore, no difference between both groups was observed in the AAV genome in the liver following the vector injection (Fig. 2E).

**Fig. 2.**
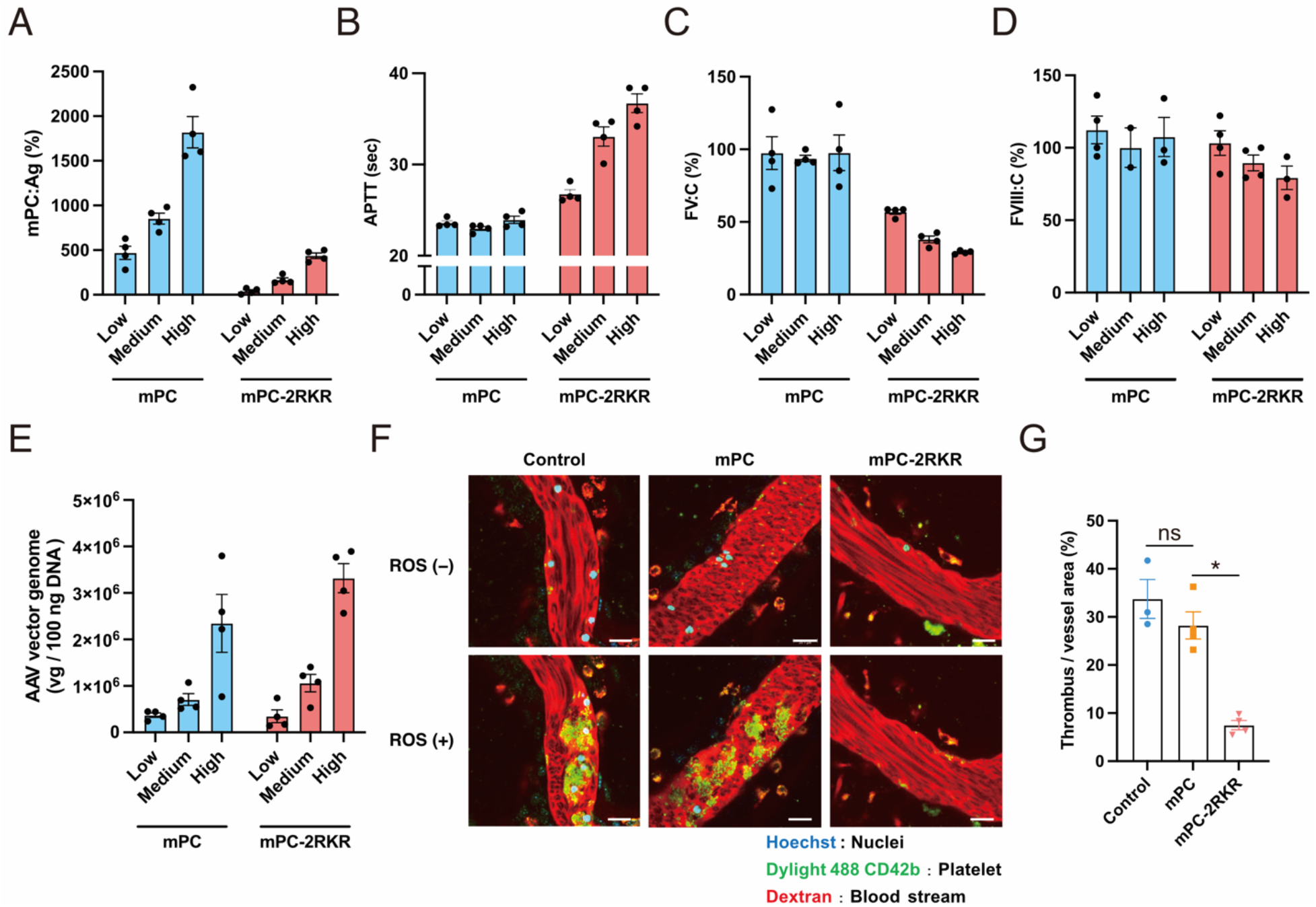
Prevention of pathogenic thrombus formation by the engineered activated protein C in mice. AAV8 vector expressing wild-type mPC or the engineered activated mPC (mPC-2RKR) under the control of HCRhAAT promoter were intravenously administered into 7-week-old C57BL/6 male mice (low, medium, and high dose: 4.0 × 10^10^, 1.2 × 10^11^, and 4.0 × 10^11^ vg/mouse, respectively). (**A**) Plasma mPC:Ag levels (mean ± SEM [n = 4]) at 4 weeks after vector injection were measured using ELISA. (**B–D**) APTT (**B**), FV:C (**C**), and FVIII:C (**D**) were measured using an automated coagulation analyzer. Values represent mean ± SEM (n = 4, except FVIII:C in mPC Medium [n = 2] and mPC-2RKR High [n = 3]). (**E**) AAV genome (mean ± SEM [n = 4]) in liver tissue at 8–12 weeks post-injection was measured using quantitative PCR. (**F**) Thrombus formation in testicular veins induced by laser-induced ROS production was observed using intravital confocal microscopy. The green signal indicates platelet thrombus formation shown by DyLight488 signals. Scale bars, 20 μm. (**G**) Signal intensities of thrombus formation were quantified using Las X Software and are expressed as the area of thrombus formation in vessel area (%). Values represent mean ± SEM (n = 3–4). \**P* < 0.05, two-tailed Student’s *t*-test. mPC, mouse protein C; HCRhAAT, an enhancer element of the hepatic control region of the Apo E/C1 gene and the human anti-trypsin promoter; mPC:Ag, mouse protein C antigen; APTT, activated partial thromboplastin time; FV:C, coagulation factor V activity; FVIII:C, coagulation factor VIII activity; ROS, reactive oxygen species; vg, vector genome; ns, not significant.

We examined the therapeutic potential of mPC-2RKR expression via AAV vector for inhibiting pathological thrombosis *in vivo*. We assessed thrombus formation in mice treated with AAV8 vector harboring mPC and mPC-2RKR at a low vector dose (4.0 × 10^10^ vg). Although the prolongation in APTT was marginal in mice treated with low-dose AAV harboring mPC-2RKR (Fig. 2B), mPC-2RKR expression significantly inhibited laser-induced ROS-elicited thrombus formation in testicular veins (Fig. 2F, G; Fig. S1; Movies S1, S3). The mPC (WT) expression did not inhibit ROS-induced thrombus formation (Fig. 2F, G; Fig. S1; Movies S1, S2).

### Phenotypic correction of PC-deficient mice via genome editing

Because conventional AAV-mediated gene therapy did not contribute to an increase in PC at the neonatal stage (Fig. S2), we employed genome editing technology to achieve a therapeutic effect during this period. We inserted the donor sequence into the *Alb* gene to target intron 14 located immediately after the terminal codon sequence *(20)*. We designed to express a target cDNA as a conjugated gene with *Alb* by the addition of a self-cleavage P2A peptide sequence instead of the terminal codon sequence of *Alb* (Fig. 3A). The target cDNA was expected to be expressed only via homology-directed repair (HDR)-mediated insertion. We confirmed that the AAV vector-mediated expression of *Staphylococcus aureus* Cas9 (SaCas9) in mice efficiently induced double-strand break (DSB) at the target site in the liver (Fig. S3A) and that it did not decrease plasma albumin levels (Fig. S3B).

**Fig. 3.**
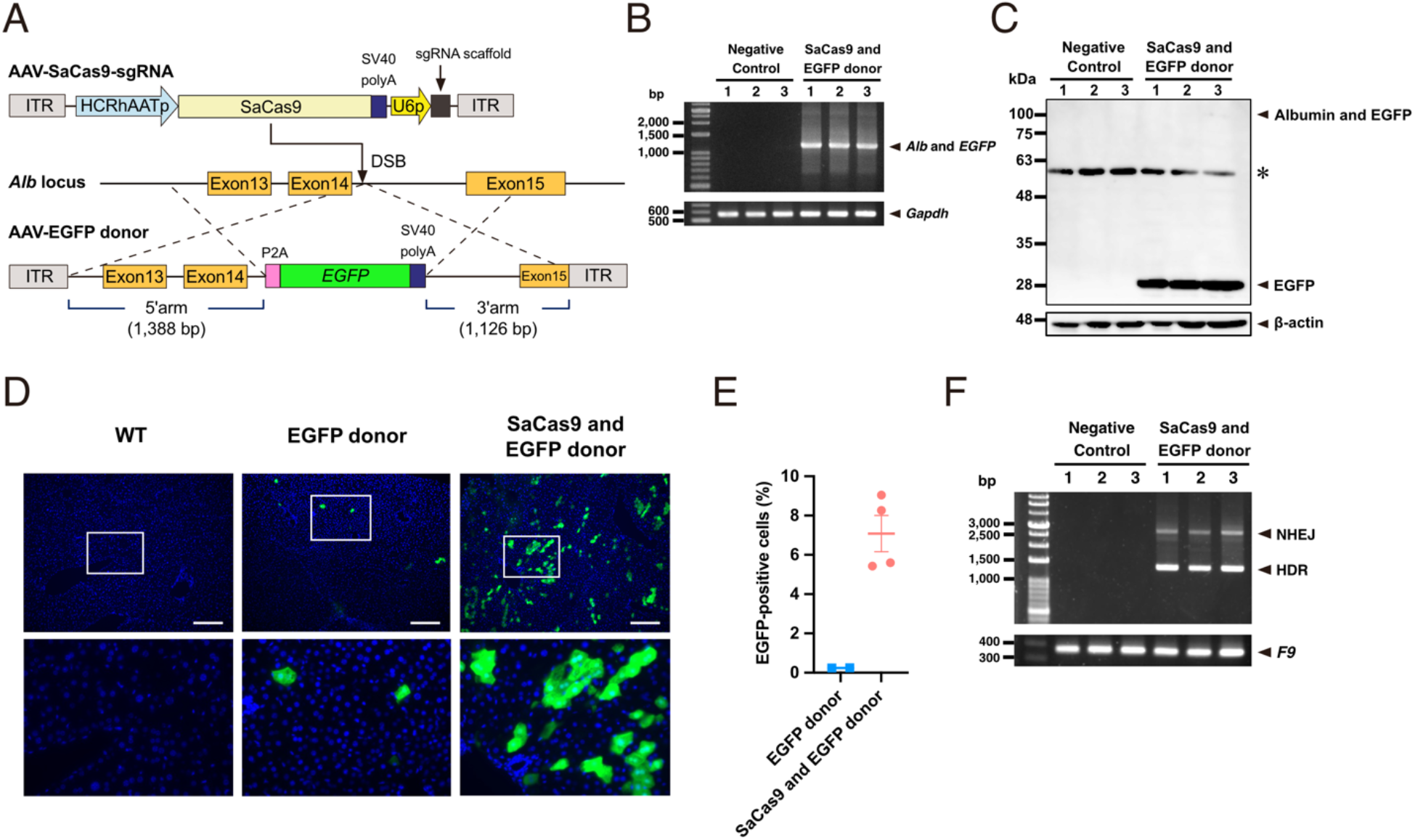
Insertion of cDNA sequence by genome editing in neonatal mice. (**A**) Schematic presentation of genome editing experiment. Self-cleaving P2A peptide sequence followed by *EGFP* cDNA was inserted into DSB induced by SaCas9 with HDR *in vivo*. (**B–G**) Neonatal C57BL/6 mice received an AAV donor vector harboring *EGFP* (2.0 × 10^11^ vg/mouse) without (EGFP donor) or with an AAV vector expressing SaCas9 and gRNA for intron 14 of *Alb* locus (6.0 × 10^10^ vg/mouse) (SaCas9 and EGFP donor). (**B**) mRNA expression of *EGFP* fused with *Alb* in the liver was detected by reverse transcription polymerase chain reaction. Representative data from three mice are shown. Liver tissues obtained from C57BL/6 were used as negative control. (**C**) Immunoblotting with anti-GFP antibody in liver tissues obtained from three mice. Representative data from three mice are shown. Liver tissues obtained from C57BL/6 were used as negative control. *A non-specific band at 60 kDa is observed in all samples. (**D**) EGFP expression in liver specimens assessed by EGFP immunostaining was photographed using an all-in-one microscope. The representative photographs were obtained from four independent experiments. Higher magnifications of the boxed regions are shown in the lower panel. Scale bars, 200 μm. (**E**) Quantitative evaluation of the EGFP-positive cells in the liver was performed with BZ-X 700 imaging software. Values represent mean ± SEM (Donor, n = 2; SaCas9 and EGFP Donor, n = 4). (**F**) PCR analysis of genomic DNA obtained from indicated organs to examine HDR and insertion at DSB by NHEJ at 6 weeks post-injection. PCR analysis can distinguish insertion via HDR and that via NHEJ by product size. DSB, double-strand break; HDR, homology-directed repair; NHEJ, non-homologous end joining; vg, vector genome.

To examine the genome editing efficacy of this system, we administered AAV8 vectors expressing SaCas9 and sgRNA (6.0 × 10^10^ vg) together with AAV8 vectors harboring P2A-fused *EGFP* cDNA sequence (2.0 × 10^11^ vg) to neonatal C57BL/6 mice (Fig. 3A). We confirmed the expression of *EGFP* mRNA conjugated with *Alb* and the efficient cleavage of EGFP protein by P2A sequence (Fig. 3B, C). Immunofluorescence staining revealed EGFP-positive cells in 7.08% ± 0.92% of all liver cells (Fig. 3D, E). Remarkably, cDNA was mainly inserted by HDR in the liver (Fig. 3F).

Further, we used mPC-2RKR for genome editing therapy to examine whether it would improve the phenotype of homozygous PC-deficient mice (*Proc*^−*/*−^ mice). We generated PC-deficient mice via genome editing and found that they died within a week (Fig. 4B). We administered same doses of AAV8 vectors expressing SaCas9 and sgRNA and AAV8 vectors harboring P2A-fused *Proc-2RKR* cDNA sequence to newborn pups born from crosses between *Proc*^*+/*−^ mice. However, all *Proc*^−*/*−^ mice died within 2–3 days after vector administration (data not shown). Because it was considered that *Proc*^−*/*−^ mice would die before achieving the efficacy of the genome editing treatment, it was necessary to ensure their survival until the appearance of therapeutic effects. We observed that crossing *Proc*^−*/*−^ mice with FVIII-deficient mice (*F8*^−*/*−^) ensured their normal survival (Fig. 4B). As previously reported *(21)*, the administration of an antibody against FVIII resulted in the complete disappearance of plasma FVIII in mice for 1 week (Fig. S5). Therefore, we periodically administered anti-FVIII antibodies to *Proc*^*+/*−^ pregnant mice, expecting that IgG would be transferred to the neonates through the placenta, following which genome editing treatment could be administered to the neonates (Fig. 4A). This method confirmed that genome editing to express mPC-2RKR ameliorated the survival of *Proc*^−*/*−^ mice (Fig. 4B). After 6 weeks of genome editing in *Proc*^−*/*−^ mice, mPC antigen levels increased by 286.5% ± 87.6% (Fig. 4C) and FV:C decreased significantly (Fig. 4D). However, the reduction of FVIII:C (Fig. 4E) and prolongation of APTT (Fig. 4F) in *Proc*^−*/*−^ mice treated with genome editing was marginal (Fig. 4E, F). Further, some *Proc*^−*/*−^ mice treated with ten times lower dose of AAV vector could also survive (Fig. 4B), but plasma FV:C was not significantly decreased (Fig. 4D). By contrast, *Proc*^−*/*−^ *F8*^−*/*−^ mice showed a significant prolongation of APTT and reduction of FVIII:C (Fig. 4E, F).

**Fig. 4.**
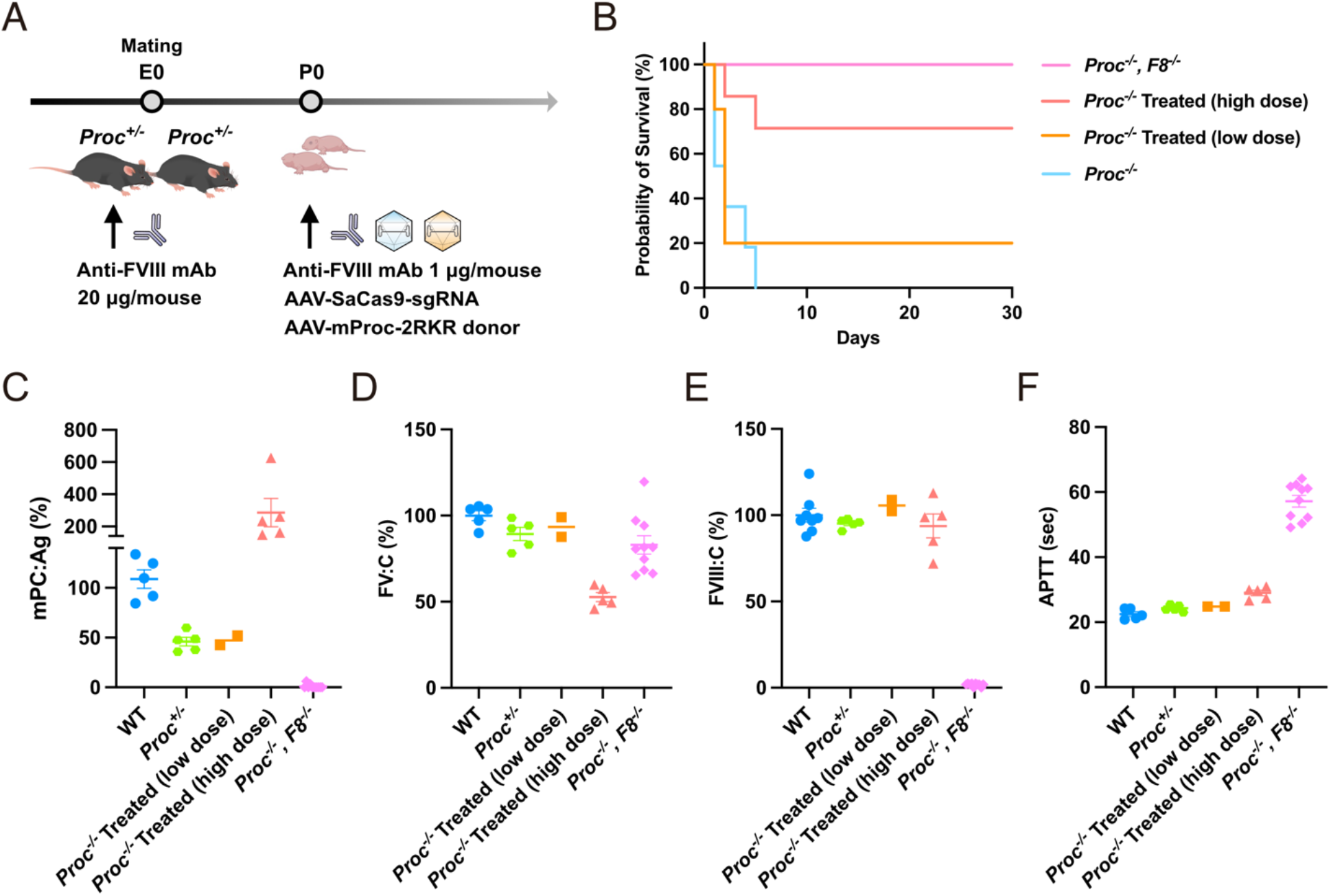
Phenotypic correction of PC-deficient homozygotic mice treated with genome editing to insert the engineered activated PC. (**A**) *Proc*^*+/*−^ female mice were treated with an FVIII antibody (#GMA8015; 20 μg/mouse) and then mated with *Proc*^*+/*−^ male mice. The antibody administration was repeated every 7 days until delivery. All neonatal mice were treated via genome editing with AAV8 vector harboring mPC-2RKR (2.0 × 10^11^ vg/mouse [high dose] or 2.0 × 10^10^ vg/mouse [low dose]) and AAV8 vector expressing SaCas9 and gRNA for intron 14 of *Alb* locus (6.0 × 10^10^ vg/mouse [high dose] or 6.0 × 10^9^ vg/mouse [low dose]), together with 1 μg of the FVIII antibody. (**B**) Kaplan– Meier survival curves for *Proc*-deficient homozygotes (*Proc*^−*/*−^, n = 13), *Proc*-deficient homozygotes treated with genome editing [*Proc*^−*/*−^ Treated (high dose), n = 9; *Proc*^−*/*−^ Treated (low dose), n = 10], and *Proc* and FVIII-double knockout mice (*Proc*^−*/*−^, *F8*^−*/*−^, n = 10). (**C–F**) The plasma mPC:Ag (**C**), FV:C (**D**), and FVIII:C (**E**), and coagulation time (APTT) (**F**). Values represent mean ± SEM (n = 5–10, except *Proc*^*-/-*^ Treated (low dose)[n = 2]). \*\*\**P* < 0.001, two-tailed Student’s *t*-test. mPC:Ag, mouse protein C antigen; APTT, activated partial thromboplastin time; FV:C, coagulation factor V activity; FVIII:C, coagulation factor VIII activity; WT, wild-type C57BL/6 mouse; *Proc*^*+/*−^, PC-deficient heterozygote; *Proc*^−*/*−^ Treated, *Proc*^−*/*−^ mice treated with genome editing; vg, vector genome.

We further assessed *in vivo* thrombus formation and bleeding tendency in *Proc*^−*/*−^ mice treated with genome editing to express mPC-2RKR. We did not observe microvascular thrombus formation in the lung and liver of *Proc*^−*/*−^ mice treated with genome editing (Fig. 5A). Compared with WT C57BL/6 mice, *Proc*^−*/*−^ mice treated with genome editing with higher vector dose and *Proc*^−*/*−^ *F8*^−*/*−^ exhibited prolonged bleeding time following tail clipping (Fig. 5C). On the other hand, bleeding after tail clipping was not significantly enhanced in *Proc*^−*/*−^ mice treated with genome editing with lower vector dose (Fig. 5C, D).

**Fig. 5.**
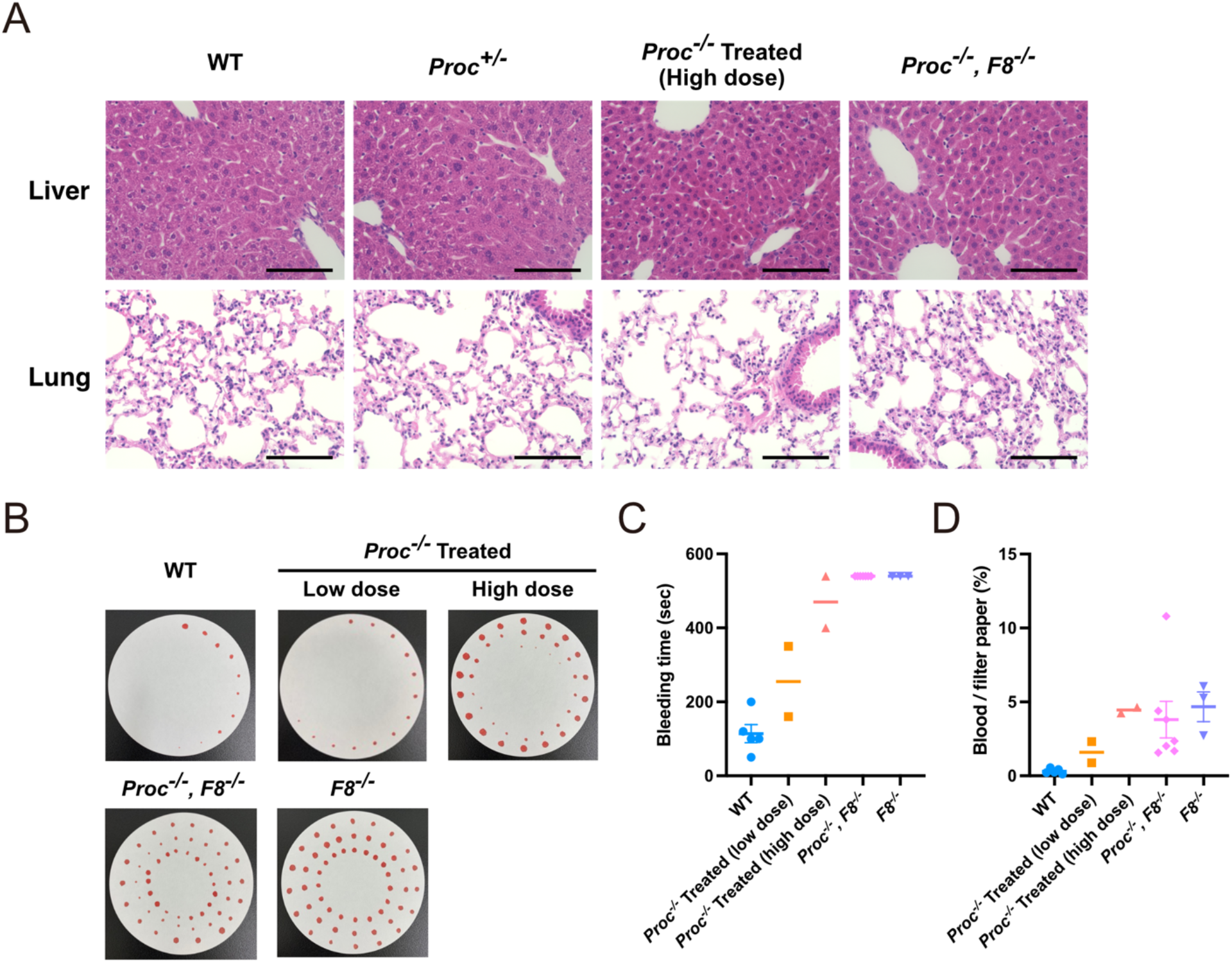
Thrombus formation and bleeding phenotype of PC-deficient homozygotic mice treated with genome editing to insert the engineered activated PC. *Proc*^*+/*−^ female mice were treated with an FVIII antibody (#GMA8015; 20 μg/mouse) and then mated with *Proc*^*+/*−^ male mice. The antibody administration was repeated every 7 days until delivery. All neonatal mice were treated via genome editing with AAV8 vector harboring mPC-2RKR (2.0 × 10^11^ vg/mouse [high dose] or 2.0 × 10^10^ vg/mouse [low dose]) and AAV8 vector expressing SaCas9 and gRNA for intron 14 of *Alb* locus (6.0 × 10^10^ vg/mouse [high dose] or 6.0 × 10^9^ vg/mouse [low dose]), together with 1 μg of the FVIII antibody. (**A**) Representative photographs of hematoxylin–eosin staining of liver and lung tissues in an indicated mouse. Scale bars,100 μm. (**B**) Representative photograph of filter paper to assess bleeding after the tail clipping in an indicated mouse. (**C, D**) Bleeding time and volume after the tail clip assay. Values represent mean ± SEM (n = 3–7, except *Proc*^*-/-*^ Treated [n = 2]). \*\*\**P* < 0.001, two-tailed Student’s *t*-test. WT, wild-type C57BL/6 mouse; *Proc*^−*/*−^ Treated, *Proc*^−*/*−^ mice treated with genome editing; *Proc*^−*/*−^, *F8*^−*/*−^, *Proc* and FVIII-double knockout mice; vg, vector genome.

## DISCUSSION

Purpura fulminans is a rare, life-threatening thrombotic disorder in newborns typically caused by homozygous or compound heterozygous abnormality in *PROC* or *PROS1 (8, 9)*. This disease generally occurs on the first day of life and rapidly progresses through multiorgan failure caused by thrombotic occlusion of vessels. Even if the patients survive with proper and timely diagnosis and treatments, neonates and children with PC or PS deficiency remain at a risk of developing severe thrombotic complications. These patients require life-long therapy with PC concentrates or fresh frozen plasma and strict anticoagulation treatments, including warfarin or heparin, under frequent monitoring *(22)*. We succeeded at expressing PC in its active form and showed that genome editing therapy using the engineered PC (PC-2RKR) at the neonatal stage could help in the survival of mice with homozygous PC deficiency. This strategy can provide a cure for patients with purpura fulminans due to PC deficiency.

We succeeded in releasing PC in its active form by inserting the furin cleavage site between the light and heavy chains. PC circulates as a zymogen, and its anticoagulant function can be achieved by releasing the activation peptide through its cleavage with thrombomodulin and thrombin complex *(23)*. The introduction of a furin cleavage site for the activation of coagulation factor VII (FVII) in gene therapy for an animal model of hemophilia has previously been reported *(24)*. Compared with the previously reported engineered PC (QGNSEDY), which enhances the binding of the Gla domain on the phospholipid surface *(18)*, PC-2RKR engineered in the present study further extended the coagulation time of human plasma. The expression of PC-2RKR in mice resulted in the prolongation of coagulation time owing to the inhibition of FV and FVIII; additionally, it led to the improvement of pathological thrombosis in WT mice and the survival of *Proc*^−*/*−^ mice.

Interestingly, although PS acts as a cofactor for efficient APC-dependent regulation of coagulation *(25)*, in the present study, even PS-deficient mice exhibited the prolongation of coagulation time by PC-2RKR. Hence, PC-2RKR may be effective for treating purpura fulminans due to PS deficiency. Moreover, compared with the WT mice, the *Proc*^−*/*−^ mice treated with genome editing to express PC-2RKR exhibited a minimal prolongation of APTT and a lower incidence of thrombus formation along with lesser bleeding time at a lower vector dose, thereby predicting that treatment with PC-2RKR led to fewer hemorrhagic complications at an appropriate dosage. Because PC-2RKR protein could be directly obtained from the cells without the cleavage of thrombin, PC-2RKR for treating purpura fulminans may be delivered as a recombinant protein or as an mRNA using lipid nanoparticle (LNP) for acute phase.

In the present study, we observed that in WT mice showing an inhibition of pathological thrombosis and *Proc*^−*/*−^ mice exhibiting an improvement of survival by PC-2RKR, albeit FV and FVIII decreased by about only 50% and 10%, respectively. In addition, *Proc*^−*/*−^ *F8*^−*/*−^ mice were viable for more than 1 year without growth retardation (data not shown), although FVIII inhibition by PC-2RKR was marginal in genome editing therapy or gene therapy. The phenotype of *Proc*^−*/*−^ *F8*^−*/*−^ mice is different from that resulting from the crossbreeding of PC knockout mice with factor XI-deficient mice *(26)*. The complete suppression of the FXI factor is insufficient to improve the phenotype, thereby leading to growth retardation and death from thrombosis within 3 months *(26)*. Currently, anticoagulation therapy for PC deficiency includes warfarin, heparin, and direct oral anticoagulants, which inhibit the production and activity of coagulation factors *(12)*. Because warfarin and heparin target multiple coagulation factors to exert anticoagulant activity, bleeding complications are major adverse events *(27)*. For PC and PS deficiency, targeting FV and FVIII, which are direct targets of PC, may provide an effective anticoagulant treatment.

The genome editing therapy targeting liver can be applied to coagulation factor deficiencies and liver metabolic diseases. Because the liver plays a central role in the metabolic processes, the abnormality of a key enzyme can progress to life-threatening conditions, such as urea cycle disorders, glycogen storage diseases, and Wilson disease *(28, 29)*. Liver transplantation remains the sole treatment strategy to cure these diseases *(30)*. Although liver transplantation has cured human PC deficiency *(31, 32)*, performing this procedure for all patients may not be possible owing to the lack of a donor or its highly invasive nature *(33)*. Gene therapy and genome editing therapy targeting the liver can become alternative therapy to cure these diseases *(20, 34, 35)*. Conventional gene therapy with AAV vectors is not expected to achieve a sustained therapeutic effect in newborns because of the potential dilution of the AAV genome during hepatocyte proliferation *(36)*.

On the other hand, the therapeutic effect of genome editing is expected to sustain for the long term, thereby leading to a disease cure. In the present study, we delivered the genome editing tool Cas9 using AAV vector; however, Cas9 expression within the cells should be transient to reduce the immunogenicity and off-target effects in the real clinical setting *(37)*. Recently, the delivery of Cas9 mRNA and gRNA with LNP has reportedly succeeded in disrupting the target gene editing in the liver for human transthyretin amyloidosis *(38)*. In the future, simultaneously administering LNP for transient Cas9 expression with an AAV vector as a donor sequence for gene insertion in the liver *in vivo* would be a better strategy.

In the current study, we inserted target genes in approximately 7% of liver cells with HDR in *in vivo* genome editing. HDR appeared to be the main mechanism for inserting the target gene because we could detect genome-edited DNA generated mainly by HDR-mediated insertion. NHEJ is the predominant repair mechanism following DSB, and the frequency of HDR is less efficient *(39)*. We have previously reported that NHEJ is the main mechanism for inserting the ectopic gene in intron 1 of the *F9* gene in FIX-deficient mice *(40)*. The efficient HDR-based insertion targeting the *Alb* locus in the present study differs from that described in our previous report, although a similar method was applied in both experiments. The only difference was that the target site and homology arms included several exon sequences in the present study. The presence of exon sequences in the arms may increase the frequency of HDRs. Further verification is warranted to determine whether HDR efficiencies differ by donor sequence in *in vivo* genome editing.

The present study has several limitations. First, the safety of performing genome editing to express the activated PC immediately after diagnosing purpura fulminans must be carefully evaluated because of the bleeding tendency owing to consumptive coagulopathy in the acute phase *(41)*. Following the replacement therapy in humans in clinical settings, it would be preferable to apply genome editing therapy after the bleeding tendency has ameliorated. Further, we must continue the protein replacement therapy for a certain period after diagnosis because manifestation of the therapeutic effects of genome editing requires several days or weeks. Under these circumstances, PC-2RKR may be applied as a protein replacement therapy or LNP preparation. In addition, the level of activated PCs to be expressed for human therapy remains unclear. In the present study, we successfully expressed a high level of activated PC in blood following genome editing. However, the minimum dosage required for the therapeutic activity should be determined before conducting a clinical trial. Finally, we only evaluated the efficacy of the genome editing treatment in a mouse model. It is necessary to verify its safety and efficacy in animal species close to humans, including non-human primates.

In conclusion, we have identified a sequence that causes the extracellular secretion of PC in its active form. We succeeded in prolonging the survival of *Proc*^−*/*−^ mice by inserting this sequence into the *Alb* locus via genome editing. The expression of PC-2RKR, but not WT PC, prevented pathological thrombosis. Even at low efficiency, modifying protein function may improve the disease phenotype. It is important to improve the efficiency and modality of genome editing and the function of the target protein to develop an effective gene therapy for intractable diseases. This study can provide a meaningful approach to gene or genome editing therapies for various diseases.

## MATERIALS AND METHODS

### Cell culture

HEK293 cells (JRCB Cell Bank, Osaka, Japan) and AAVpro 293T cells (Takara Bio Inc., Shiga, Japan) were cultured in Dulbecco’s modified Eagle Medium (Sigma Aldrich, Saint Louis, MO) supplemented with 10% fetal bovine serum (Thermo Fisher Scientific, Waltham, MA) and 2 mM L-glutamine (Thermo Fisher Scientific).

### Plasmid constructs and AAV vector production

The cDNA encoding hPC was synthesized by GenScript Japan Inc. (Tokyo, Japan). The cDNA of mPC was amplified via reverse transcription polymerase chain reaction using total RNA from the mouse liver as a template. Total RNA was prepared using an RNA purification kit (RNeasy Mini kit; Qiagen Inc., Valencia, CA). The isolated RNA was reverse transcribed using the PrimeScript™ RT reagent kit (Takara Bio Inc.). Reverse-transcribed cDNA was amplified using PrimeStar DNA polymerase (Takara Bio Inc.) and cloned into pCR-Blunt II-TOPO^®^ (Thermo Fisher Scientific). cDNAs were incorporated into pcDNA3 (Thermo Fisher Scientific). The inserted sequences were synthesized by GenScript Japan Inc. to insert self-cleaving sites between the light chain and heavy chain of PC, following which these sequences were inserted into the parental plasmid using In-Fusion^®^ cloning kit (Takara Bio Inc.).

A plasmid comprising a chimeric promoter (HCRhAAT, which is an enhancer element of the hepatic control region of the Apo E/C1 gene and the human anti-trypsin promoter), cDNAs, and the SV40 polyadenylation signal was created to specifically express target gene in hepatocytes by AAV vector. The DNA fragment was introduced between inverted terminal repeats into the pAAV plasmid to produce the AAV vector. sgRNA sequences were designed using online software provided by Benchling (https://benchling.com) (Table S1). For the production of AAV vector that would induce DSB, a DNA fragment comprising a chimeric HCRhAAT promoter, SaCas9 cDNA, the SV40 polyadenylation signal, and a sgRNA sequence driven by the U6 promoter was introduced between inverted terminal repeats into the pAAV plasmid. We simultaneously created a pAAV donor plasmid containing a P2A self-cleaving peptide sequence-conjugated cDNA sequence possessing1.0 kb homologous arms at the target site for inserting the target gene via HDR.

The AAV genes were packaged by triple plasmid transfection of AAVpro293T cells to generate the AAV vector (helper-free system), as described previously *(42)*. pHelper plasmid (Takara Bio Inc.), the plasmid expressing Rep and serotype 8 capsid (AAV8), and the gene transfer plasmid were simultaneously transfected. AAV vectors were purified from the transfected cells after 72 h using the ultracentrifugation method, as previously described *(43)*. The titration of recombinant AAV vectors was performed using quantitative PCR, as previously described *(40)*.

### Plasmid transfection of the cells with wild or engineered PC

The plasmid was transfected into HEK293 cells using Lipofectamine 3000 Reagent (Thermo Fisher Scientific) according to the manufacturer’s instructions. To establish a stable cell line, linearized pcDNA3 plasmid expressing the cDNA of WT or engineered PC was transfected to HEK293 cells. To select the transfected cell clones, G418 (Nacalai Tesque Inc., Kyoto, Japan) was added to the culture medium to aid the selection. To measure PC in the supernatant, 5 μg/mL of vitamin K (menatetrenone, Eisai Inc., Tokyo, Japan) was supplemented into the culture medium.

### PC, FV, and FVIII activities, coagulation time, and ELISA

The activity of hPC (hPC:C) was measured using Berichrom PROTEIN C (Sysmex, Kobe, Japan) by an automated coagulation analyzer (CS-1600; Sysmex). When measuring the existence of APC in the sample, a PC activator (snake venom to activate hPC) was not added to samples. The activities of FV (FV:C) and FVIII (FVIII:C) were measured using a one-stage clotting-time assay on an automated coagulation analyzer (CS-1600; Sysmex). PT and APTT were measured by an automated coagulation analyzer (CA-500; Sysmex) using Thrombocheck PT and Thrombocheck APTT (Sysmex). hPC antigen (hPC:Ag) was measured using the Human Protein C AssayMaxTM ELISA Kit (Assaypro, St. Charles, MO) according to the manufacturer’s instructions. Mouse plasma albumin was measured by an automated analyzer (7180 Clinical Analyzer; Hitachi, Tokyo, Japan) at Oriental Yeast Co. Ltd (Tokyo, Japan). To measure mPC:Ag, 96-well plates were coated with anti-mouse protein C polyclonal sheep IgG (R&D Systems, Minneapolis, MN) and incubated at 4°C overnight, following which they were blocked with phosphate-buffered saline (PBS) containing 5% casein (EMD Millipore Corp., Billerica, MA). Mouse plasma samples were added into wells and incubated for 1 h. After extensive washing with PBS-T, anti-mouse protein C polyclonal sheep IgG conjugated with horse-radish peroxidase (GeneTex, Irvine, CA) was incubated for 1 h. Antibody binding was visualized using ABTS^®^ Peroxidase Substrate (KPL Protein Research Products, WA). The optical density of each well was measured at 415 nm.

### Animal experimentation

All animal experimental procedures were approved by The Institutional Animal Care and Concern Committee of Jichi Medical University (permission number: 20080-04), and animal care was conducted in accordance with the committee’s guidelines and ARRIVE guidelines. FVIII-deficient mice (B6;129S4-*F8*^*tm1Kaz*^/J) were kindly provided by Dr. H. H. Kazazian Jr. (University of Pennsylvania, Philadelphia, PA). C57BL/6J male mice were purchased from Japan SLC (Shizuoka, Japan). Animals were maintained in isolators in the specific pathogen-free facility of Jichi Medical University at 23°C ± 3°C with a 12:12 h light/dark cycle.

To obtain plasma samples, mice were anesthetized with isoflurane (1–3%), and the blood sample was drawn from the jugular vein using a 29G micro-syringe (TERUMO Corp., Tokyo, Japan) containing 1/10 (volume/volume) sodium citrate. Platelet-poor plasma was isolated by centrifugation and then frozen and stored at −80°C until the analysis. AAV vector was administered intravenously through the jugular vein (100–150 μL) and intraperitoneally (10 μL) in adult and neonatal mice, respectively. When indicated, an anti-FVIII monoclonal antibody (#GMA8015, Green Mountain Antibodies, Burlington, VT) was intraperitoneally administered (20 μg/body in adult mice and 1 μg/body in neonatal mice).

The designed sgRNAs for the PC-deficient mice were purchased from Integrated DNA Technologies Inc. (Coralville, IA; Table S1). To generate PC-deficient mice, a mixture of *Streptococcus pyogenes* Cas9 protein (100 ng/μL; Integrated DNA Technologies Inc.) and an sgRNA (3 μmol) was electroporated into embryos (20–100 cells) using a NEPA21 Type II Super Electroporator (Nepa Gene, Chiba, Japan). The zygotes were cultured until the two-cell stage, following which they were transferred into pseudopregnant female mice. The sgRNA sequence is described in Table S1.

### Intravital microscopy

Intravital microscopy was performed as reported previously *(40, 44)*. Briefly, anesthetized mice were injected with anti-platelet GPIbβ conjugated with DyLight488 (0.01 mg/body; Emfret Analytics GmbH & Co., Eibelstadt, Germany), rhodamine B isothiocyanate (5 mg/body; Sigma Aldrich) or Texas Red-Dextran (2.5 mg/body; Thermo Fisher Scientific), and Hoechst 33342 (2 mg/body; Thermo Fisher Scientific); hematoporphyrin (0.125 mg/body Sigma Aldrich, Germany) was injected to produce ROS on laser irradiation. Blood cell dynamics and thrombus formation at the testicular vein were visualized using laser excitation and ROS production (wavelength 488 nm, 1.5 mW power) at a 63× objective lens. Sequential images of the testicular vein (40–60 μm) were obtained using a confocal microscope (Leica TCS SP8, Leica Microsystems, Wetzlar, Germany) with a high-sensitive GsAsP hybrid detector. The signal intensity of thrombus formation (shown by DyLight488 signals) was quantified using Las X Software (Leica Microsystems).

### T7 endonuclease assay

Genomic mutations induced by DSB were detected using the T7 endonuclease assay. PCR fragments were amplified with ExTaq DNA polymerase (Takara Bio Inc.). Purified PCR products were denatured and re-annealed using a thermal cycler, following which they were treated with T7 endonuclease (Nippon Gene, Tokyo, Japan). DNA fragments were analyzed using agarose gel electrophoresis or microchip electrophoresis system (MCE-202 MultiNA; Simazu Corp., Tokyo, Japan).

### Quantification of AAV in the genome and mRNA expression

Mice deeply anesthetized with isoflurane were perfused with 50 mL PBS to remove the blood, and then tissues were isolated. Genomic DNA and RNA were extracted using DNeasy Blood & Tissue Kit (Qiagen Inc.) and RNeasy Mini Kit (Qiagen Inc.), respectively. The RNA samples were reverse transcribed using a PrimeScript RT Reagent kit (Takara Bio Inc.). Quantitative real-time PCR was performed using THUNDERBIRD™ Probe qPCR Mix or THUNDERBIRD™ SYBR qPCR Mix (TOYOBO, Osaka, Japan) with a QuantStudio™ 12K Flex Real-Time PCR system (Thermo Fisher Scientific). The amount of the AAV genomic DNA was measured by quantifying the SV40 polyadenylation signal. The ectopic expression levels of PC mRNA were normalized to mRNA levels of *Hprt1*. The primer and probe sequences are described in Table S1.

### Immunoblotting

Liver tissue was lysed using RIPA buffer (50 mM Tris [pH 7.5], 150 mM NaCl, 0.1% SDS, 1% NP-40, 0.5% sodium deoxycholate, cOmplete^®^ Mini Protease Inhibitor Cocktail [Roche Diagnosis, Basel, Switzerland], 10 mM NaF, and 1 mM Na_3_VO_4_), and sonicated in a Beads Crusher μT-12 (TAITEC Corp., Saitama, Japan). The lysate was centrifuged at 15,000 ×*g* for 10 min. The supernatant was recovered as the total liver lysate. Total liver lysates were resolved using sodium dodecyl sulfate–polyacrylamide gel electrophoresis and then transferred to a polyvinylidene difluoride membrane. The membranes were blocked using 5% (w/v) skimmed milk powder in TBS-T buffer (20 mM Tris [pH 7.5], 150 mM NaCl, 0.05% Tween 20) for 1 h. After extensive washing with TBS-T, the membranes were incubated overnight with anti-GFP polyclonal antibody (MBL Co., Aichi, Japan) in TBS-T containing 5% (w/v) bovine serum albumin at 4°C. Antibody binding was detected using HRP-conjugated anti-rabbit IgG and visualized with Immobilon Western Chemiluminescent HRP Substrate (Millipore, Burlington, MA) and an ImageQuant LAS4000 digital imaging system (GE Healthcare, Buckinghamshire, UK).

### Histological analysis and immunohistochemistry

Mice anesthetized with isoflurane were perfused with 50 mL PBS. The tissues were fixed in 10% formalin, embedded in paraffin or with 4% paraformaldehyde, incubated with PBS containing sucrose (10%–30%), and frozen in the presence of optimal cutting temperature compound (Sakura Fintek Japan, Tokyo, Japan). Following this, the paraffin-embedded tissue sections were dewaxed in xylene, rehydrated with ethanol, washed with water, and processed for hematoxylin and eosin staining. For immunohistochemistry, tissue sections were blocked with 5% donkey serum and then incubated with an anti-GFP polyclonal antibody (MBL Co.). The sections were then incubated with anti-rabbit IgG conjugated with AlexaFluor 488 (Thermo Fisher Scientific) for 2 h at 4°C. Slides were mounted using VECTASHIELD Mounting Medium with DAPI (Vector Laboratories, Burlingame, CA, USA). The results of hematoxylin and eosin staining and immunofluorescence staining were observed and photographed using an all-in-one microscope (BZ-X700; Keyence, Tokyo, Japan). The quantitative evaluation of EGFP-positive cells was performed using BZ-X 700 imaging software (Keyence).

### Tail clip assay

The tail clip assay was performed to quantify the bleeding tendency. The tail of the mice anesthetized with isoflurane was clipped with a scalpel. The injured tail was placed on filter paper every 10 sec. Bleeding time was assessed as the time to stop bleeding. Bleeding volume was quantified as the percentage of the red bleeding area in the filter paper.

### Statistical analyses

Statistical analyses were performed using GraphPad Prism^®^ (GraphPad Software, San Diego, CA). All data are presented as the mean ± standard error of the mean (SEM) values. The method for statistics in each experiment is described in Figure Legends.

## SUPPLEMENTARY MATERIALS

Table S1. sgRNA and the oligonucleotide primer pairs used in this study.

Fig. S1. Prevention of pathogenic thrombus by the engineered activated protein C in mice.

Fig. S2. Ectopic expression of hPC by AAV vector.

Fig. S3. Genome editing at exon 14 of *Alb* locus located immediately after terminal codon.

Fig. S4. Generation of *Proc-*deficient mice.

Fig. S5. Depletion of plasma FVIII by antibody administration in mice.

Fig. S6. Whole gel images of Fig. 3.

Movie S1. Thrombus formation in the testicular vein in C57BL/6 mouse.

Movie S2. Thrombus formation in the testicular vein in C57BL/6 mouse treated with an AAV vector expressing wild-type mPC.

Movie S3. Thrombus formation in the testicular vein in C57BL/6 mouse treated with an AAV vector expressing the engineered mPC-2RKR.

## Acknowledgments

We acknowledge Dr. H.H. Kazazian Jr. (University of Pennsylvania, Philadelphia, PA) for providing *F8*-deficient mice. We thank Yaeko Suto, Mika Kishimoto, Tamaki Aoki, Sachiyo Kamimura, Mai Hayashi, Yuiko Ogihara, Nagako Sekiya, Tomoko Noguchi, Hiromi Ozaki, and Hiroko Hayakawa of Jichi Medical University for their technical assistance. Optima XE-90 was subsidized by JKA through its promotion funds from KEIRIN RACE.

## Author contributions

Conceptualization: TT, TO

Methodology: TT, NB, YN, YK, ON, TO

Investigation: TT, NB, YN, YK, MH, NK, TH, TO

Visualization: NB, TF

Funding acquisition: ON, TO

Project administration: ON, TO

Supervision: EM, ON, TO

Writing – original draft: TT, TO

Writing – review & editing: TT, NB, YN, YK, MH, NK, TH, TF, EM, ON, TO

## Data and materials availability

The original data and materials within the paper are available from the corresponding author upon request. *Proc*-deficient mice have been deposited at Riken BRC (#RBRC11381, Ibaraki, Japan). Materials transfer agreements are required for the donation of plasmids and mice.

## REFERENCES AND NOTES

1. A. M. Wendelboe, G. E. Raskob, Global Burden of Thrombosis: Epidemiologic Aspects. Circ Res 118, 1340–1347 (2016).

2. E. Previtali, P. Bucciarelli, S. M. Passamonti, I. Martinelli, Risk factors for venous and arterial thrombosis. Blood Transfus 9, 120–138 (2011).

3. P. Bucciarelli, S. M. Passamonti, E. Biguzzi, F. Gianniello, F. Franchi, P. M. Mannucci, I. Martinelli, Low borderline plasma levels of antithrombin, protein C and protein S are risk factors for venous thromboembolism: Anticoagulant proteins and venous thromboembolism. Journal of Thrombosis and Haemostasis 10, 1783–1791 (2012).

4. A. R. Rezaie, H. Giri, Anticoagulant and signaling functions of antithrombin. J Thromb Haemost 18, 3142–3153 (2020).

5. J. H. Griffin, J. A. Fernández, A. J. Gale, L. O. Mosnier, Activated protein C: Activated protein C. Journal of Thrombosis and Haemostasis 5, 73–80 (2007).

6. C. T. Esmon, The protein C anticoagulant pathway. Arterioscler Thromb 12, 135–145 (1992).

7. B. Lipe, D. L. Ornstein, Deficiencies of Natural Anticoagulants, Protein C, Protein S, and Antithrombin. Circulation 124 (2011), doi:10.1161/CIRCULATIONAHA.111.044412.

8. M. J. Manco-Johnson, T. C. Abshire, L. J. Jacobson, R. A. Marlar, Severe neonatal protein C deficiency: Prevalence and thrombotic risk. The Journal of Pediatrics 119, 793–798 (1991).

9. C. Mahasandana, V. Suvatte, RichardA. Marlar, MarilynJ. Manco-Johnson, LindaJ. Jacobson, WilliamE. Hathaway, Neonatal purpura fulminans associated with homozygous protein S deficiency. The Lancet 335, 61–62 (1990).

10. P. Knoebl, Severe congenital protein C deficiency: the use of protein C concentrates (human) as replacement therapy for life-threatening blood-clotting complications. BTT, 285 (2008).

11. P. Monagle, A. K. C. Chan, N. A. Goldenberg, R. N. Ichord, J. M. Journeycake, U. Nowak-Göttl, S. K. Vesely, Antithrombotic Therapy in Neonates and Children. Chest 141, e737S–e801S (2012).

12. P. Monagle, C. A. Cuello, C. Augustine, M. Bonduel, L. R. Brandão, T. Capman, A. K. C. Chan, S. Hanson, C. Male, J. Meerpohl, F. Newall, S. H. O’Brien, L. Raffini, H. van Ommen, J. Wiernikowski, S. Williams, M. Bhatt, J. J. Riva, Y. Roldan, N. Schwab, R. A. Mustafa, S. K. Vesely, American Society of Hematology 2018 Guidelines for management of venous thromboembolism: treatment of pediatric venous thromboembolism. Blood Advances 2, 3292–3316 (2018).

13. N. A. Goldenberg, M. J. Manco-Johnson, Protein C deficiency: PROTEIN C DEFICIENCY. Haemophilia 14, 1214–1221 (2008).

14. A. Thompson, Structure, function, and molecular defects of factor IX. Blood 67, 565–572 (1986).

15. L. A. George, S. K. Sullivan, A. Giermasz, J. E. J. Rasko, B. J. Samelson-Jones, J. Ducore, A. Cuker, L. M. Sullivan, S. Majumdar, J. Teitel, C. E. McGuinn, M. V. Ragni, A. Y. Luk, D. Hui, J. F. Wright, Y. Chen, Y. Liu, K. Wachtel, A. Winters, S. Tiefenbacher, V. R. Arruda, J. C. M. van der Loo, O. Zelenaia, D. Takefman, M. E. Carr, L. B. Couto, X. M. Anguela, K. A. High, Hemophilia B Gene Therapy with a High-Specific-Activity Factor IX Variant. N Engl J Med 377, 2215–2227 (2017).

16. P. Chowdary, S. Shapiro, M. Makris, G. Evans, S. Boyce, K. Talks, G. Dolan, U. Reiss, M. Phillips, A. Riddell, M. R. Peralta, M. Quaye, D. W. Patch, E. Tuddenham, A. Dane, M. Watissée, A. Long, A. Nathwani, Phase 1–2 Trial of AAVS3 Gene Therapy in Patients with Hemophilia B. N Engl J Med 387, 237–247 (2022).

17. A. Gruber, J. Griffin, Direct detection of activated protein C in blood from human subjects. Blood 79, 2340–2348 (1992).

18. Y.-H. Sun, L. Shen, B. Dahlbäck, Gla domain–mutated human protein C exhibiting enhanced anticoagulant activity and increased phospholipid binding. Blood 101, 2277–2284 (2003).

19. P. M. Gempeler-Messina, K. Volz, B. Bühler, C. Müller, Protein C activators from snake venoms and their diagnostic use. Haemostasis 31, 266–272 (2001).

20. A. De Caneva, F. Porro, G. Bortolussi, R. Sola, M. Lisjak, A. Barzel, M. Giacca, M. A. Kay, K. Vlahoviček, L. Zentilin, A. F. Muro, Coupling AAV-mediated promoterless gene targeting to SaCas9 nuclease to efficiently correct liver metabolic diseases. JCI Insight 4, e128863 (2019).

21. S. Keshava, J. Sundaram, A. Rajulapati, U. R. Pendurthi, L. V. M. Rao, Pharmacological concentrations of recombinant factor VIIa restore hemostasis independent of tissue factor in antibody-induced hemophilia mice. J Thromb Haemost 14, 546–550 (2016).

22. A. Minford, L. R. Brandão, M. Othman, C. Male, R. Abdul-Kadir, P. Monagle, A. D. Mumford, D. Adcock, B. Dahlbäck, P. Miljic, M. T. DeSancho, J. Teruya, Diagnosis and management of severe congenital protein C deficiency (SCPCD): Communication from the SSC of the ISTH. J of Thrombosis Haemost 20, 1735–1743 (2022).

23. C. T. Esmon, The Protein C Pathway. Chest 124, 26S–32S (2003).

24. P. Margaritis, V. R. Arruda, M. Aljamali, R. M. Camire, A. Schlachterman, K. A. High, Novel therapeutic approach for hemophilia using gene delivery of an engineered secreted activated Factor VII. J. Clin. Invest. 113, 1025–1031 (2004).

25. M. Gierula, J. Ahnström, Anticoagulant protein S—New insights on interactions and functions. J. Thromb. Haemost. 18, 2801–2811 (2020).

26. J. C. Y. Chan, J. G. Ganopolsky, I. Cornelissen, M. A. Suckow, M. J. Sandoval-Cooper, E. C. Brown, F. Noria, D. Gailani, E. D. Rosen, V. A. Ploplis, F. J. Castellino, The Characterization of Mice with a Targeted Combined Deficiency of Protein C and Factor XI. The American Journal of Pathology 158, 469–479 (2001).

27. S. Schulman, R. J. Beyth, C. Kearon, M. N. Levine, Hemorrhagic complications of anticoagulant and thrombolytic treatment: American College of Chest Physicians Evidence-Based Clinical Practice Guidelines (8th Edition). Chest 133, 257S–298S (2008).

28. M. Odievre, Clinical presentation of metabolic liver disease. J Inherit Metab Dis 14, 526–530 (1991).

29. T. Taddei, P. Mistry, M. L. Schilsky, Inherited metabolic disease of the liver. Curr Opin Gastroenterol 24, 278–286 (2008).

30. G. Mazariegos, B. Shneider, B. Burton, I. J. Fox, N. Hadzic, P. Kishnani, D. H. Morton, S. Mcintire, R. J. Sokol, M. Summar, D. White, V. Chavanon, J. Vockley, Liver transplantation for pediatric metabolic disease. Molecular Genetics and Metabolism 111, 418–427 (2014).

31. M. J. Lee, K. M. Kim, J. S. Kim, Y. J. Kim, Y. J. Lee, T. T. Ghim, Long-term survival of a child with homozygous protein C deficiency successfully treated with living donor liver transplantation. Pediatric Transplantation 13, 251–254 (2009).

32. M. Matsunami, A. Ishiguro, A. Fukuda, K. Sasaki, H. Uchida, T. Shigeta, H. Kanazawa, S. Sakamoto, M. Ohta, H. Nakadate, R. Horikawa, A. Nakazawa, M. Ishige, K. Mizuta, M. Kasahara, Successful living domino liver transplantation in a child with protein C deficiency. Pediatr Transplantation 19, E70–E74 (2015).

33. A. S. Bodzin, T. B. Baker, Liver Transplantation Today: Where We Are Now and Where We Are Going: Review Articles. Liver Transpl 24, 1470–1475 (2018).

34. R. Sharma, X. M. Anguela, Y. Doyon, T. Wechsler, R. C. DeKelver, S. Sproul, D. E. Paschon, J. C. Miller, R. J. Davidson, D. Shivak, S. Zhou, J. Rieders, P. D. Gregory, M. C. Holmes, E. J. Rebar, K. A. High, In vivo genome editing of the albumin locus as a platform for protein replacement therapy. Blood 126, 1777–1784 (2015).

35. Y. Yang, L. Wang, P. Bell, D. McMenamin, Z. He, J. White, H. Yu, C. Xu, H. Morizono, K. Musunuru, M. L. Batshaw, J. M. Wilson, A dual AAV system enables the Cas9-mediated correction of a metabolic liver disease in newborn mice. Nat Biotechnol 34, 334–338 (2016).

36. S. C. Cunningham, A. Spinoulas, K. H. Carpenter, B. Wilcken, P. W. Kuchel, I. E. Alexander, AAV2/8-mediated Correction of OTC Deficiency Is Robust in Adult but Not Neonatal Spfash Mice. Molecular Therapy 17, 1340–1346 (2009).

37. E. A. Taha, J. Lee, A. Hotta, Delivery of CRISPR-Cas tools for in vivo genome editing therapy: Trends and challenges. Journal of Controlled Release 342, 345–361 (2022).

38. J. D. Gillmore, E. Gane, J. Taubel, J. Kao, M. Fontana, M. L. Maitland, J. Seitzer, D. O’Connell, K. R. Walsh, K. Wood, J. Phillips, Y. Xu, A. Amaral, A. P. Boyd, J. E. Cehelsky, M. D. McKee, A. Schiermeier, O. Harari, A. Murphy, C. A. Kyratsous, B. Zambrowicz, R. Soltys, D. E. Gutstein, J. Leonard, L. Sepp-Lorenzino, D. Lebwohl, CRISPR-Cas9 In Vivo Gene Editing for Transthyretin Amyloidosis. N Engl J Med 385, 493–502 (2021).

39. S. V. Vartak, S. C. Raghavan, Inhibition of nonhomologous end joining to increase the specificity of CRISPR/Cas9 genome editing. FEBS J 282, 4289–4294 (2015).

40. T. Ohmori, Y. Nagao, H. Mizukami, A. Sakata, S. Muramatsu, K. Ozawa, S. Tominaga, Y. Hanazono, S. Nishimura, O. Nureki, Y. Sakata, CRISPR/Cas9-mediated genome editing via postnatal administration of AAV vector cures haemophilia B mice. Sci Rep 7, 4159 (2017).

41. J.-F. Dhainaut, S. B. Yan, Y.-E. Claessens, Protein C/activated protein C pathway: Overview of clinical trial results in severe sepsis: Critical Care Medicine 32, S194–S201 (2004).

42. J. Mimuro, H. Mizukami, S. Hishikawa, T. Ikemoto, A. Ishiwata, A. Sakata, T. Ohmori, S. Madoiwa, F. Ono, K. Ozawa, Y. Sakata, Minimizing the Inhibitory Effect of Neutralizing Antibody for Efficient Gene Expression in the Liver With Adeno-associated Virus 8 Vectors. Molecular Therapy 21, 318–323 (2013).

43. N. Baatartsogt, Y. Kashiwakura, M. Hayakawa, N. Kamoshita, T. Hiramoto, H. Mizukami, T. Ohmori, A sensitive and reproducible cell-based assay via secNanoLuc to detect neutralizing antibody against adeno-associated virus vector capsid. Molecular Therapy - Methods & Clinical Development 22, 162–171 (2021).

44. S. Nishimura, I. Manabe, M. Nagasaki, S. Kakuta, Y. Iwakura, N. Takayama, J. Ooehara, M. Otsu, A. Kamiya, B. G. Petrich, T. Urano, T. Kadono, S. Sato, A. Aiba, H. Yamashita, S. Sugiura, T. Kadowaki, H. Nakauchi, K. Eto, R. Nagai, In vivo imaging visualizes discoid platelet aggregations without endothelium disruption and implicates contribution of inflammatory cytokine and integrin signaling. Blood 119, e45–e56 (2012).

