## Supplemental Table 1, Figure 1-6 for "Cure of congenital purpura fulminans via expression of engineered protein C through neonatal genome editing in mice"

**Affiliations:**
[1]Department of Clinical Laboratory Science, Division of Health Sciences, Graduate School of Medical Science, Kanazawa University; Ishikawa, 920-0942, Japan.
[2]Department of Biochemistry, Jichi Medical University School of Medicine; Tochigi, 329-0498, Japan.
[3]Center for Experimental Medicine, Jichi Medical University; Tochigi, 329-0498, Japan.
[4]Center for Gene Therapy Research, Jichi Medical University; Tochigi, 329-0498, Japan.
[5]Division of Cell and Molecular Medicine, Center for Molecular Medicine, Jichi Medical University; Tochigi, 329-0498, Japan.
[6]Department of Cardiovascular Medicine, The University of Tokyo Hospital; Tokyo, 113-8655, Japan.
[7]Department of Biological Sciences, Graduate School of Science, The University of Tokyo; Tokyo, 113-0033, Japan.

### SUPPLEMENTARY MATERIALS

Table S1. sgRNA and the oligonucleotide primer pairs used in this study.

Fig. S1. Prevention of pathogenic thrombus by the engineered activated protein C in mice.

Fig. S2. Ectopic expression of hPC by AAV vector.

Fig. S3. Genome editing at exon 14 of *Alb* locus located immediately after terminal codon.

Fig. S4. Generation of *Proc*-deficient mice.

Fig. S5. Depletion of plasma FVIII by antibody administration in mice.

Fig. S6. Whole gel images of Fig. 3.

Table S1. sgRNA and the oligonucleotide primer pairs used in this study.

Movie S1. Thrombus formation in the testicular vein in C57BL/6 mouse.

Movie S2. Thrombus formation in the testicular vein in C57BL/6 mouse treated with an AAV vector expressing the wild-type mPC.

Movie S3. Thrombus formation in the testicular vein in C57BL/6 mouse treated with an AAV vector expressing the engineered mPC-2RKR.

**Table S1.** sgRNA and the oligonucleotide primer pairs used in this study.

| Target gene | Cas9 | sgRNA | Sequence | PAM sequence |
|---|---|---|---|---|
| *Alb* Intron 14 | *Staphylococcus aureus* | | GATGACCATACGTGAAGACCT | AAGAGT |
| *Proc* Exon 9 | *Streptococcus pyogenes* | sgRNA1 | TCGTCCACCCTAACTACACC | CGG |
| | | sgRNA2 | CACTCATTTCGAGCAACCAA | AGG |

| Target gene | | Sequence |
|---|---|---|
| T7 endonuclease assay | | |
| *Alb* Intron 14 | F | 5'-GCCACACTGCTGCCTATTAAATACC-3' |
| | R | 5'-GGACTCCACATAGTGGTTCATGTAAG-3' |
| *Proc* Exon 9 | F | 5'-ATTCTTCCTAGCAGTGCCCT-3' |
| | R | 5'-CCCAGCCTTACTCCAAGAGT-3' |
| Real time qPCR | | |
| *PROC* mRNA | F | 5'-GCCAACTCCTTCCTGGAGGA-3' |
| | R | 5'-TCCTTGGCCTCCTCGAAGTC-3' |
| *Hprt1* mRNA | F | 5'-GTTGGATACAGGCCAGACTTTGTTG-3' |
| | R | 5'-GATTCAACTTGCGCTCATCTTAGGC-3' |
| SV40 poly A | F | 5'-AGCAATAGCATCACAAATTTCACAA-3' |
| | R | 5'-CCAGACATGATAAGATACATTGATGAGTT-3' |
| | probe | 5'-AGCATTTTTTTCACTGCATTCTAGTTGTGGTTTGTC-3' |
| Detection of HDR and NHEJ | | |
| *Alb-EGFP* mRNA | F | 5'-CTCAGGTGTCAACCCCAACT-3' |
| | R | 5'-AGGACCATGTGATCGCGCTTCT-3' |
| *Gapdh* mRNA | F | 5'-CCATCACCATCTTCCAGGAG-3' |
| | R | 5'-CCTGCTTCACCACCTTCTTG-3' |
| *EGFP* knock into *Alb* Intron 14 | F | 5'-CGTCATGGGTGTGACTTTTG-3' |
| | R | 5'-TTGCTCACAGGTCCAGGGTTCTCCTCCA-3' |
| mouse *F9* Exon 3 and Intron 3 | F | 5'-TGGAAGCAGTATGTTGGTAAGC-3' |
| (internal control) | R | 5'-AACAGGGATAGTAAGATTGTTCC-3' |

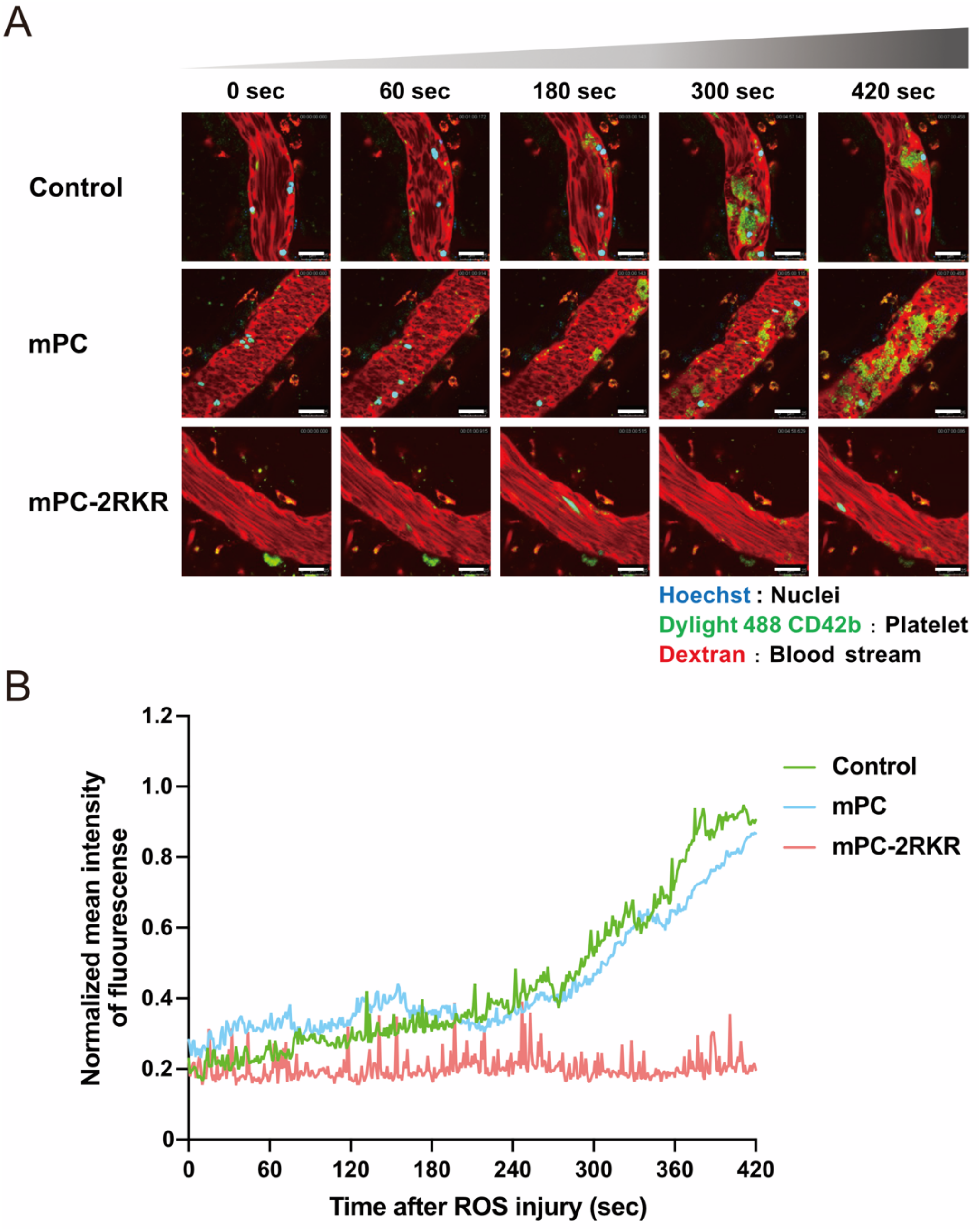


**Fig. S1. Prevention of pathogenic thrombus by the engineered activated protein C in mice.** AAV8 vector expressing wild-type mPC or the engineered activated mPC (mPC-2RKR) under the control of HCRhAAT promoter were intravenously administrated into 7-week-old C57BL/6 male mice ($4.0 \times 10^{10}$ vg/mouse). (**A**) Thrombus formation in testicular veins induced by laser-elicited ROS production was observed using intravital confocal microscopy (Leica TCS SP8, Leica Microsystems). Green signal indicates platelet thrombus formation shown by DyLight488 signals. Scale bars, 25 µm. (**B**) Representative signal intensities of thrombus formation were quantified using Las X Software (Leica Microsystems). mPC, mouse protein C; HCRhAAT, an enhancer element of the hepatic control region of the Apo E/C1 gene and the human anti-trypsin promoter; ROS, reactive oxygen species; vg, vector genome.

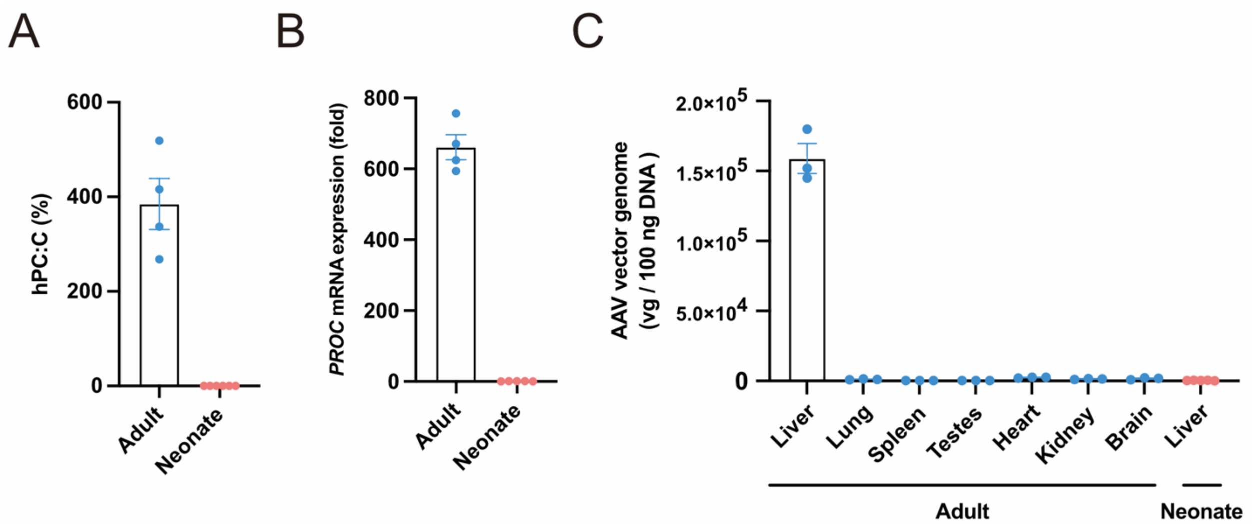


**Fig. S2**. **Ectopic expression of hPC by AAV vector.** AAV8 vector expressing wild-type hPC under the control of HCRhAAT promoter was administrated into 7-week-old (adult) ($4.0 \times 10^{10}$ vg/mouse) or neonatal C57BL/6 mice ($2.0 \times 10^{9}$ vg/mouse). (**A**) Plasma levels of hPC:C were measured at 8 weeks after vector administration. Values represent mean ± SEM (n = 4–6). (**B**) mRNA expression of *PROC* in the liver was measured using real-time reverse transcription polymerase chain reaction and was expressed as the fold increase in the *PROC*/*Hprt1* ratio. Values represent mean ± SEM (n = 4–5). (**C**) Distribution of the AAV genome in each tissue at 8 weeks after the injection in mice treated at 7 weeks of age or neonatal mice. Values represent mean ± SEM (n = 3–5). HCRhAAT, an enhancer element of the hepatic control region of the Apo E/C1 gene and the human anti-trypsin promoter; hPC:C, human protein C activity; SEM, standard error of the mean; vg, vector genome.

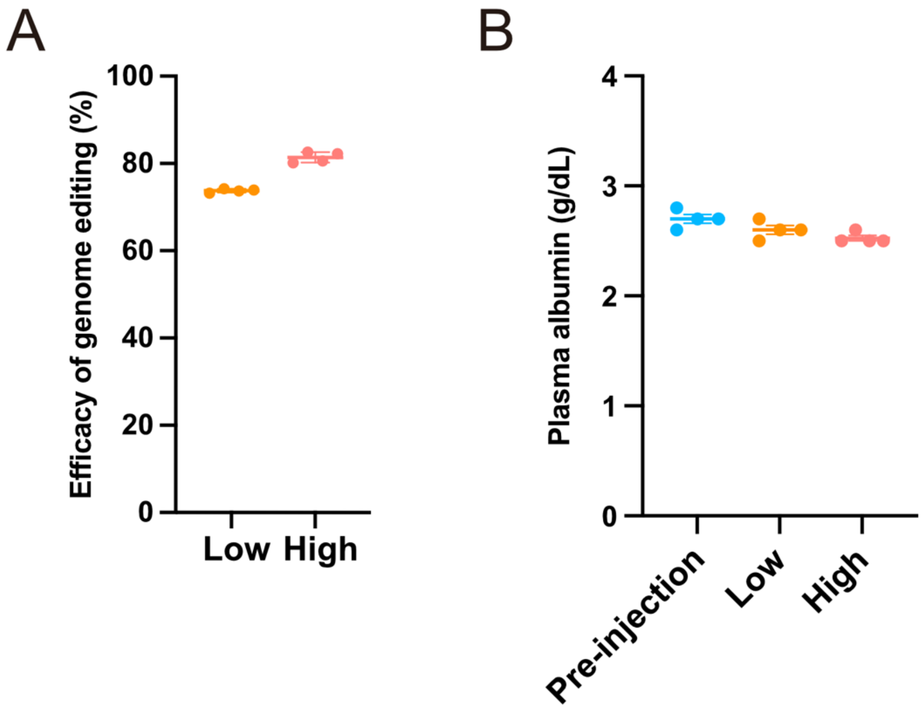


**Fig. S3. Genome editing at exon 14 of *Alb* locus located immediately after terminal codon.** Adult C57BL/6 mice were treated with an AAV8 vector expressing SaCas9 and sgRNA8 targeting *Alb* at $1.0 \times 10^{11}$ vg/mouse (Low) or $3.0 \times 10^{11}$ vg/mouse (High). (**A**) SaCas9-mediated double-strand break (DSB) at *Proc* in the liver at 8 weeks after the vector injection was assessed using T7 endonuclease assay. Values represent mean ± SEM (n = 4). (**B**) Plasma levels of albumin before (Pre-injection) and after the vector injection (8 weeks) were assessed by ELISA. Values represent mean ± SEM (n = 4). vg, vector genome.

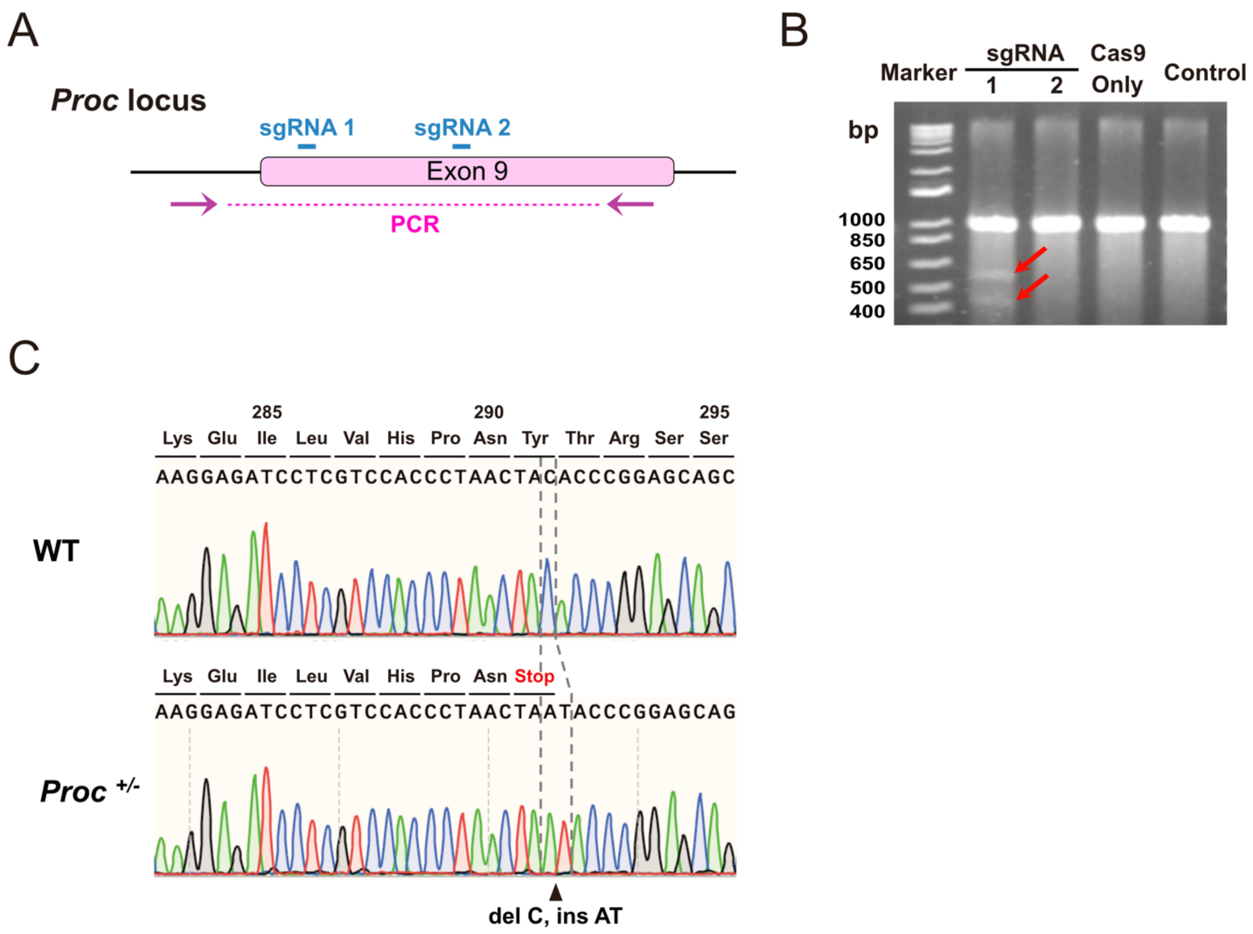


**Fig. S4. Generation of *Proc*-deficient mice.** (**A**) Two sgRNA sequences were designed at exon 9 of the *Proc* locus. The location of PCR primers is described using red arrows. (**B**) SpCas9-mediated double-strand break at *Proc* in TLR-3 cells was assessed using T7 endonuclease assay. TLR-3 cells were transduced without (Control) or with a plasmid expressing SaCas9 and an indicated gRNA [without gRNA (Cas9 only), gRNA1, gRNA2] targeting the exon 9 of *Proc*. Red arrows represent a cleaved DNA by T7 endonuclease. (**C**) DNA sequence of allele obtained from wild-type or heterozygote of *Proc*-deficient mice created by genome editing.

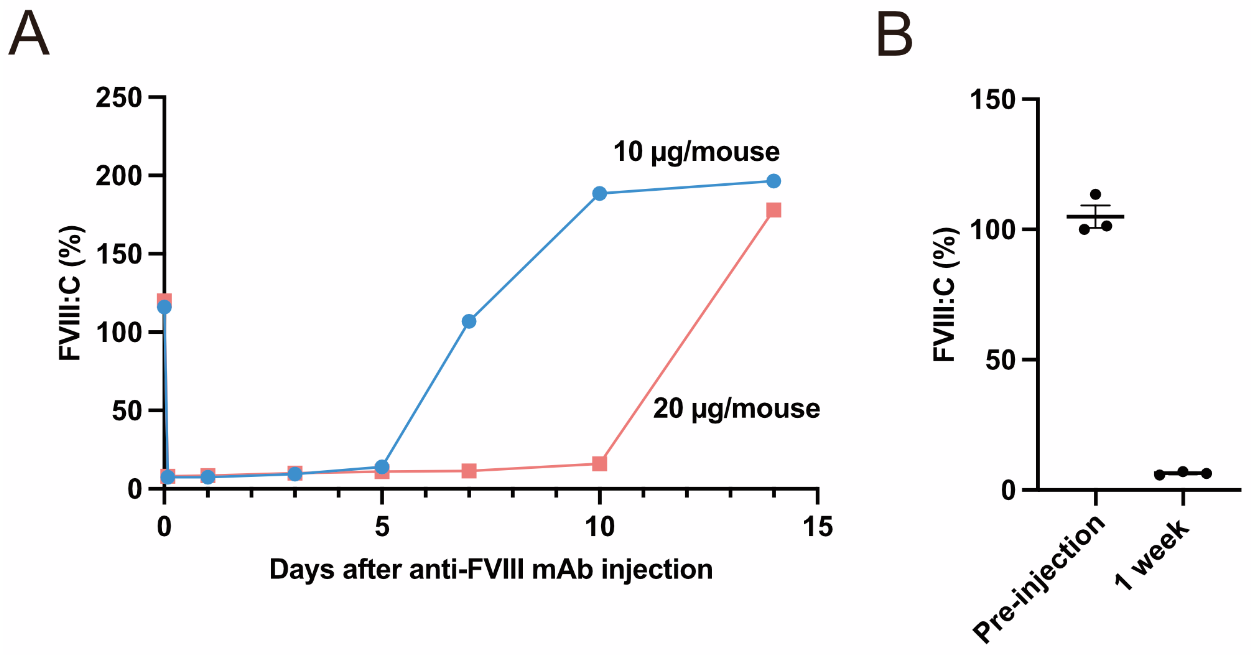


**Fig. S5. Depletion of plasma FVIII by antibody administration in mice.** (**A**) C57BL/6 mice (7-week-old males) were intraperitoneally treated with 10 µg or 20 µg anti-FVIII antibody (#GMA8015, Green Mountain Antibodies). Blood was obtained at indicated times after the antibody injection. Plasma levels of FVIII activity (FVIII:C) were measured by one clotting assay. Each data set is representative of one mouse. (**B**) Pregnant heterozygotes of *Proc*-deficient mice were treated with the 20 µg anti-FVIII antibody. Plasma FVIII:C at 0 (Pre-injection) and 7 days after the antibody injection. Values represent mean ± SEM (n = 3).

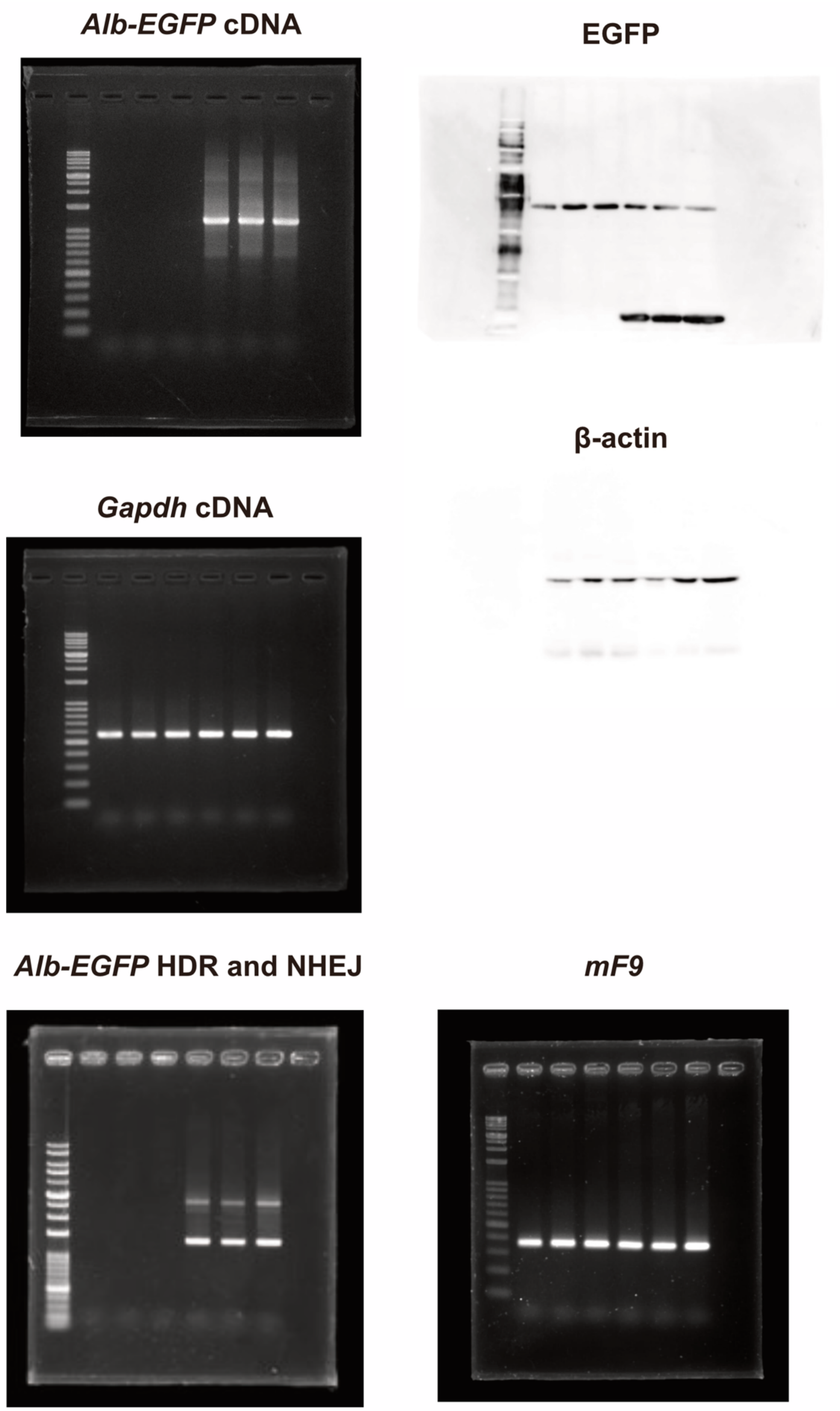


**Fig. S6. Whole gel images of Fig. 3.**
